# Cardiomyocyte PGC-1α enables physiological adaptations to endurance exercise through suppression of GDF15 and cardiac atrophy

**DOI:** 10.1101/2024.01.30.578093

**Authors:** Sumeet A. Khetarpal, Haobo Li, Tevis Vitale, James Rhee, Louisa Grauvogel, Claire Castro, Melanie J. Mittenbühler, Nicholas E. Houstis, Ariana Vargas-Castillo, Amanda L. Smythers, Jing Liu, Casie Curtin, Hans-Georg Sprenger, Katherine A. Blackmore, Alexandra Kuznetsov, Rebecca Freeman, Dina Bogoslavski, Patrick T. Ellinor, Aarti Asnani, Phillip A. Dumesic, Pere Puigserver, Jason D. Roh, Bruce M. Spiegelman, Anthony Rosenzweig

**Affiliations:** Corrigan Minehan Heart Center and Cardiovascular Research Center, Massachusetts General Hospital, Boston, MA, USA; Department of Cancer Biology, Dana Farber Cancer Institute, Boston, MA, USA; Department of Cell Biology, Harvard Medical School, Boston, MA, USA; Department of Anesthesia, Critical Care, and Pain Medicine, Massachusetts General Hospital, Boston, MA, USA; CardioVascular Institute, Beth Israel Deaconess Medical Center, Boston, MA, USA; Cardiovascular Disease Initiative, the Broad Institute of Harvard and MIT; Institute for Heart and Brain Health, University of Michigan, Ann Arbor, MI, USA

**Keywords:** Exercise, Endurance training, Peroxisome proliferator activated receptor coactivator 1 alpha (PGC-1α), Growth differentiation factor 15 (GDF15), Cardiac atrophy, Cardiac fibrosis, Senescence associated secretory phenotype (SASP)

## Abstract

Exercise training induces physiological cardiac hypertrophy, enhanced mitochondrial biogenesis and myocardial contractility. In skeletal muscle, the transcriptional coactivator PGC-1α is a key orchestrator of these responses. The heart expresses abundant and exercise-responsive PGC-1α, but it is unclear whether cardiomyocyte PGC-1α is necessary for cardiac adaptation to endurance training. Here we demonstrate that cardiomyocyte PGC-1α is required for physiological cardiac hypertrophy during exercise training in mice. In the absence of cardiomyocyte PGC-1α, voluntary wheel running does not improve exercise capacity and instead confers immune-fibrotic-atrophic heart failure after just 6 weeks of training. We identify cardiomyocyte PGC-1α as a negative regulator of stress-responsive senescence gene expression. The most enriched of these is the myomitokine GDF15. GDF15 is secreted locally but not systemically in PGC-1α-deficient mouse hearts and reduces cardiomyocyte size. Cardiomyocyte-specific reduction of GDF15 expression preserves exercise tolerance and cardiac contractility in PGC-1α-deficient mice during endurance training. Finally, we show that cardiomyocyte *PPARGC1A* expression correlates with cardiomyocyte number and negatively with GDF15 expression in human cardiomyopathies through single nucleus RNA sequencing. Our data implicate cardiomyocyte PGC-1α as a vital safeguard against stress-induced atrophy and local GDF15-induced dysfunction during exercise.

## Introduction

Endurance exercise confers remarkable protection from the incidence and adverse sequelae of cardio-metabolic and other chronic diseases^1–4^. Endurance training induces myriad adaptations including intra- and inter-organ communication through protein secretion, mitochondrial function, detoxification of circulating compounds, and local and systemic inflammatory modulation^5–9^.

Response to endurance training is best understood in the muscle. Here exercise training induces multiple adaptations including mitochondrial biogenesis, myokine protein and metabolite secretion, muscle hypertrophy and others^10^. A vital orchestrator of these responses is the transcriptional co-activator peroxisome proliferator activated receptor 1α (PGC-1α)^11,12^. Tissue-specific gain- and loss-of-function studies have established its necessity and sufficiency for the optimal training response^13,14^. The heart is a mitochondrial- and PGC-1α-rich muscle that must also adapt to exercise training. The role of PGC-1α in mitochondrial biogenesis and oxidative metabolism in the heart has been described in studies of disease models in mice lacking PGC-1α globally^10,11,14–16^. However, whether cardiomyocyte PGC-1α is needed for benefits in the adaptive stress of endurance exercise training is unclear.

Here, we investigated the role of cardiomyocyte PGC-1α in acute exercise and endurance training through genetic loss-of-function in mice. We find that cardiomyocyte PGC-1α is required for the beneficial cardiac response to exercise training in mice. Its absence causes exercise-induced immune fibrotic heart failure, cardiac atrophy and accelerated age-related myocardial gene expression including local induction of the myomitokine GDF15. Silencing cardiomyocyte GDF15 mitigates heart failure and exercise intolerance in vivo.

## Results

### Cardiomyocyte PGC-1α is required to adapt to endurance training in mice

To investigate whether cardiomyocyte PGC-1α is required for exercise tolerance, we studied young adult (12-week-old) male WT and cardiomyocyte PGC-1α-deficient (KO) mice. Consistent with prior studies of these mice, sedentary KO mice (KO_Sed) demonstrated normal heart dimensions and contractility at rest by echocardiography (**Figure 1A**)^16^. They did exhibit a ∼13% reduction in resting heart rate compared to the WT sedentary (WT_Sed) group (**Figure 1A**). While cardiomyocyte PGC-1α deficiency did not impair resting contractility, we observed that heart *Ppargc1a* gene expression increased acutely 4-5 fold in the heart within 30 minutes of exhaustive treadmill exercise in WT mice (**Figure S1**). We thus tested acute exhaustive treadmill exercise tolerance in WT and KO mice. Sedentary KO mice demonstrated similar work capacity at maximum effort as wild-type littermates, though they achieved approximately 12% less maximum speed (p<0.0001) and 15% less maximum distance ran at exhaustion (p<0.001) (**Figure 1B**). Despite modest reduction in peak exercise tolerance, KO mice demonstrated a 25% decrease in their contractility measured as fractional shortening (FS) at peak stress compared to WT mice (decreased contractile reserve), which expectedly augmented their FS with exercise (p<0.0001, **Figure 1C-D**).

**Figure 1:**
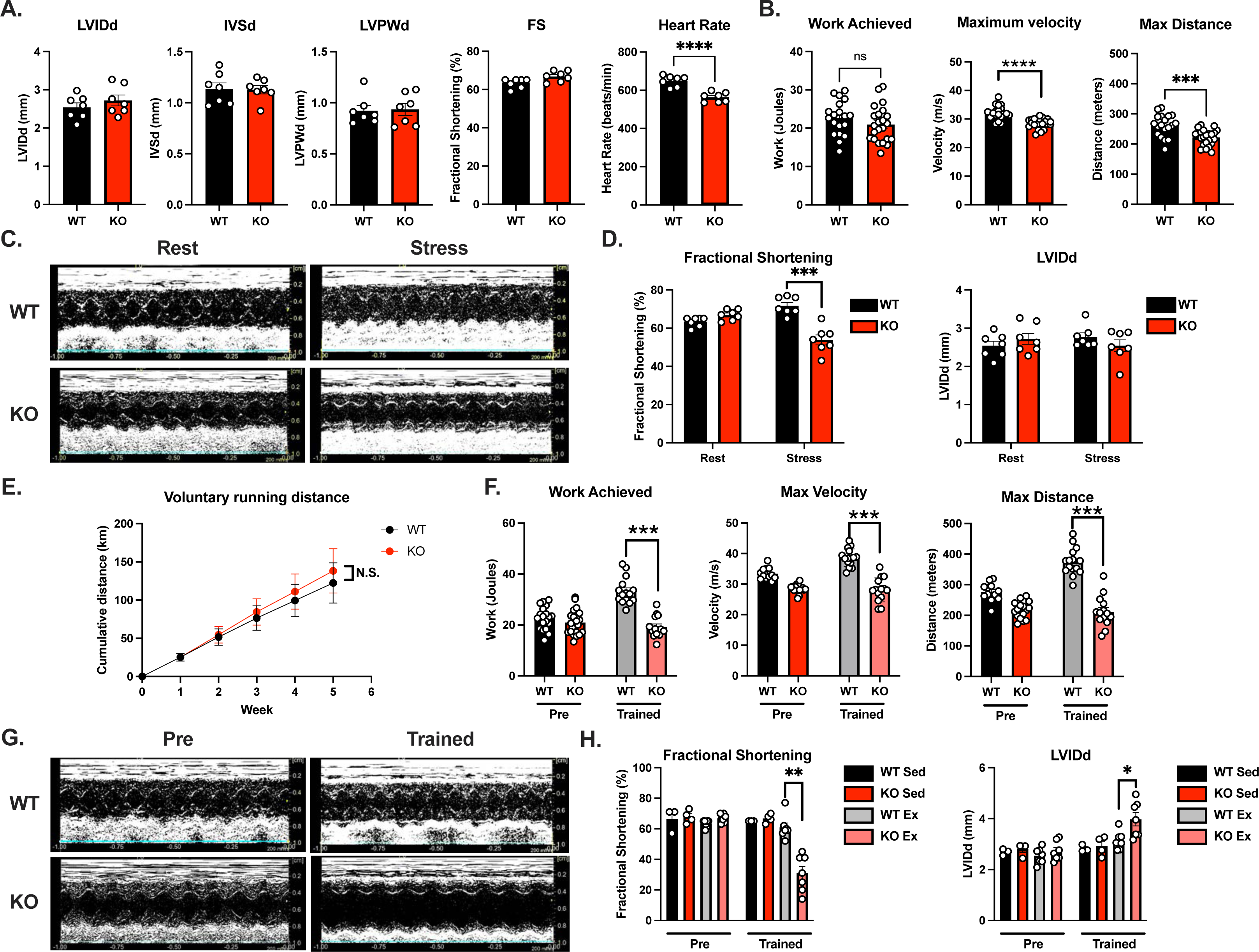
Failure to adapt to endurance exercise training in cardiomyocyte PGC-1⍺ deficient mice. **A**. Conscious murine echocardiography measures of left ventricular internal diameter in diastole (LVIDd), interventricular septum dimension in diastole (IVSd), left ventricular posterior wall thickness in diastole (LVPWd), fractional shortening (FS), and heart rate (HR) in sedentary WT or KO mice. **B**. Acute treadmill exercise test results for work output achieved, maximum velocity, and max distance run at exhaustion in mice from A. **C**. M-mode murine echocardiographic images of mice in B at rest and immediately after cessation of treadmill running. **D.** (left) FS and LVIDd at rest and with peak stress in mice from B. (right) **E**. Cumulative voluntary running distance by WT and KO mice over 5 weeks of endurance exercise training. **F.** Acute treadmill exercise testing in mice from E before and after 6 weeks of voluntary wheel running. **G.** Resting M-mode murine echocardiographic images of mice in E-F before training and at 6 weeks of endurance training. **H.** FS and LVIDd from rest echocardiographic images from G both before and after 6 weeks of wheel running. Data is expressed as mean +/- S.E.M.. For A and B, ***P<0.001, ****P<0.0001, Unpaired T-test; for D-H, *P<0.05, **P<0.01, ***P<0.001, Unpaired T-test with Welch’s correction.

We hypothesized that endurance training could augment exercise capacity in cardiac PGC-1α-deficient mice. This was based on inference from models of PGC-1α-deficiency in other tissues^13,18^. Additionally, we previously showed that endurance training in mice induced as many as 175 transcriptional programs associated with physiological cardiac hypertrophy, suggesting many potential programs aside from PGC-1α that could confer exercise adaptation^19^. To test this, we trained 12-week-old WT and KO mice through voluntary wheel running for up to 8 weeks. We assessed exercise tolerance by treadmill testing prior to and at 6 weeks of voluntary wheel running. WT and KO mice voluntarily ran comparable distances over the training period (**Figure 1E**). After 6 weeks of training, WT mice (WT_Ex) demonstrated a ∼50% increase in their exercise tolerance during treadmill testing , but KO mice failed to demonstrate any improvement (**Figure 1F**). Surprisingly, KO mice after exercise training (KO_Ex) now demonstrated a ∼53% *reduction* in resting FS (p<0.001, **Figure 1G-H**) and dilation of the left ventricle with a ∼46% increase in left ventricular cavity size (p<0.05, **Figure 1G-H**).

### Immune-fibrotic heart failure in exercise-trained cardiomyocyte PGC-1α deficiency

We explored the unexpected quick onset of contractile dysfunction with exercise training in the KO_Ex mice. At 8 weeks of training, these mice demonstrated systemic features of heart failure. This included increased heart and lung weight (**Figure 2A**) and pathologic cardiac gene expression (Nppb, Myh7 to Myh6 ratio; **Figure 2B**). The KO_Ex mice also demonstrated reduced inguinal white adipose tissue (iWAT) and gastrocnemius weights and increased iWAT oxidative gene expression, consistent with early systemic wasting (**Figure S2**). Given PGC-1α’s known central role in promoting mitochondrial biogenesis^20^, we quantified relative heart mitochondrial mass by measuring the relative amount of mtDNA to nuclear DNA in the hearts of our mice. WT_Ex mice demonstrated increased relative mtDNA relative to WT_Sed mice whereas KO_Ex mice showed no increase (**Figure 2C**). RNA sequencing revealed a strong geneset enrichment^21,22^ of hallmark genes comprising mitochondrial oxidative phosphorylation (OXPHOS^23^) in WT mice after endurance training. In the KO mice there was a relative depletion of this same geneset after exercise training (**Figure 2D-E**). Comparing heart transcriptomes of sedentary and exercise trained mice, KO mice demonstrated increased representation^24^ of genes important in cell cycle progression and decreased expression of genes related to calcium ion transport particularly in the sedentary state. After exercise training, KO mice demonstrated increased expression of genes involved in immunocyte chemotaxis, protein secretion and extracellular matrix deposition and a relative deficiency in oxidative gene expression (**Figure 2F-G**). Cell type estimation from RNA seq data using murine and human immunocyte marker-based deconvolution suggested an enrichment for macrophages in the KO_Ex hearts (**Figure 2G and Figure S3A-B**). KO_Ex hearts showed increased pro-inflammatory CD68+ monocyte/macrophage infiltration in their myocardium (**Figure 2H**). They also showed increased CD3+ cells consistent with an overall increase in cardiac inflammation (**Figure S3C**). KO_Ex hearts contained increased collagen fibrosis measured by Picrosirius Red staining (**Figure 2I**) and increased pro-fibrotic protein expression including TGFβ and periostin (**Figure S3D**). These findings imply the development of immune-fibrotic heart failure in the KO mice after exercise training.

**Figure 2:**
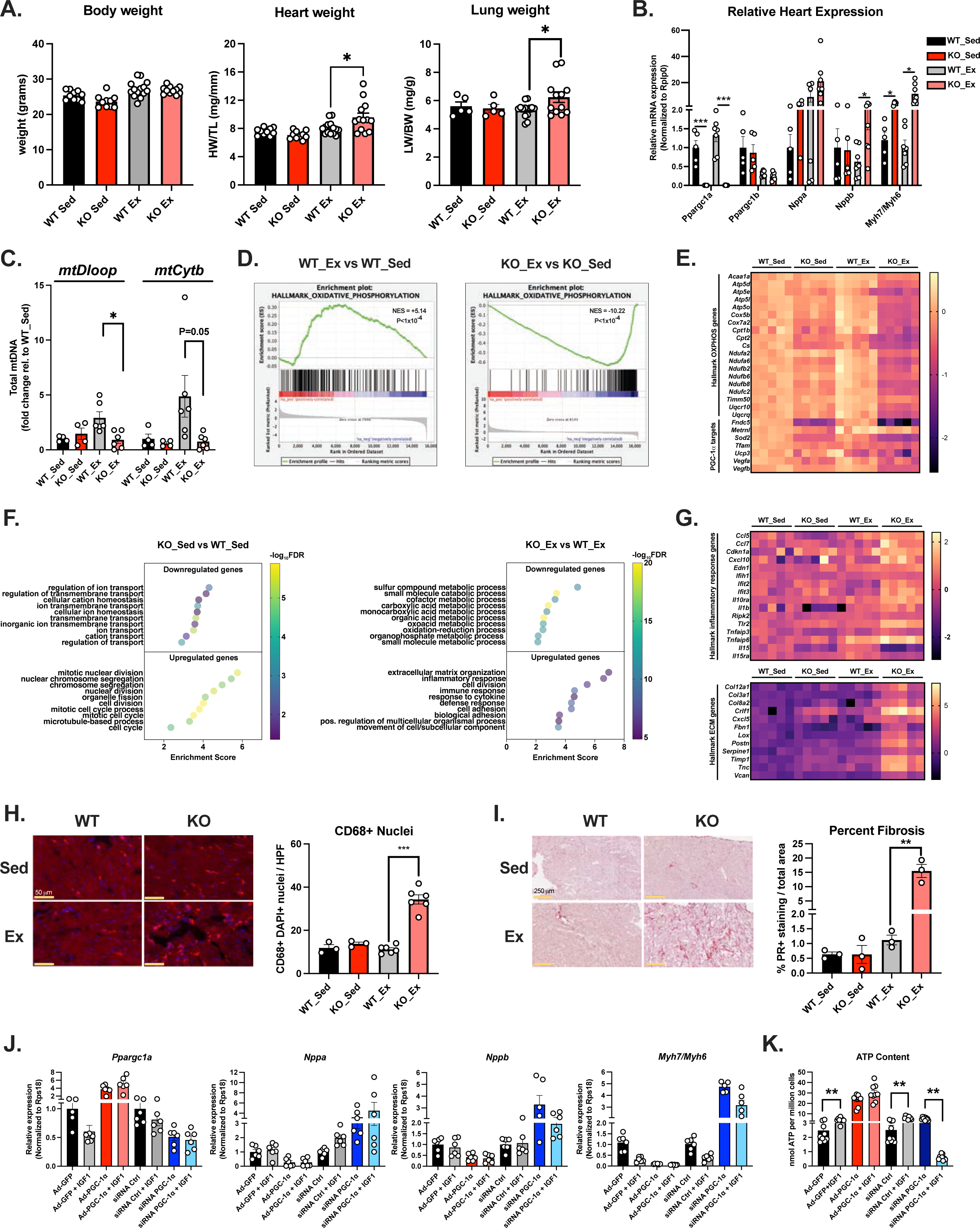
Immune-fibrotic heart failure in exercise-trained cardiomyocyte PGC-1⍺ KO mice. **A**. Body weight, heart weight relative to tibia length (HW/TL) and lung weight normalized to body weight in sedentary (Sed) or 8 week exercise trained (Ex) WT or KO mice. **B**. Bulk heart gene expression by quantitative RT-PCR from the mice in A. Expression of indicated genes was normalized to that of Rplp0. **C**. Mitochondrial DNA (mtDNA) in total heart DNA extracts from mice in A. Total mtDNA was measured by quantitative RT-PCR using mtDNA primers mtDloop and mtCytb and normalized for each sample to nuclear DNA copy number using primers for β-actin. **D**. Gene set enrichment plot for Hallmark OXPHOS genes in the comparison of WT_Ex vs WT_Sed mice (left) and KO_Ex vs KO_Sed mice (right) using bulk heart RNA seq normalized gene expression. **E**. Heatmap of relative gene expression from RNA seq data expressed as log_2_(fold change) relative to the WT_Sed group for representative hallmark OXPHOS genes and selected known PGC-1α targets. **F**. Top 10 Gene Ontology Biological Process terms for downregulated (top) and upregulated (bottom) genes (Padj < 0.05) from RNA seq data in D for comparison of KO_Sed vs WT_Sed groups (left) and KO_Ex vs WT_Ex groups (right). Enrichment scores are plotted as dots with color corresponding to enrichment score FDR value according to the colorscale. **G**. Heatmap of relative gene expression from RNA seq data for representative Hallmark inflammatory response genes (top) and Hallmark extracellular matrix (ECM) genes (bottom). **H**. Representative images from CD68 (pink), PCM1 (red), and DAPI (blue) staining of frozen heart sections from indicated mice (left) and quantification of CD68+/DAPI+ stains per high power field (HFP) from mice of each group (right). **I**. Representative images from Picrosirius red collagen staining from indicated mice (left) and quantification of positive staining per HPF (right). **J**. Relative mRNA expression of indicated genes in NRVMs treated with recombinant adenovirus expressing GFP (Ad-GFP), PGC-1α (Ad-PGC-1α), siRNA to a scramble control sequence (siCtrl) or to rat PGC-1α (siPGC-1α), and with or without IGF1 treatment for 48 hrs prior to cell lysis for RNA extraction. Expression was normalized to that of Rps18. **K**. ATP concentration in NRVM cell lysates. Data is expressed as mean +/- S.E.M.. *P<0.05, **P<0.01, ***P<0.001, **** P<0.0001, student’s unpaired T-test with Welch’s correction.

To further investigate the mechanism of exercise-induced dysfunction in the KO_Ex mice, we studied neonatal rat ventricular myocytes (NRVMs) in culture^25^. Adenoviral overexpression of PGC-1α increased transcript levels ∼6-fold and reduced stress-related gene expression (Nppa, Nppb, Myh7/Myh6) (**Figure 2J**). siRNA mediated PGC-1α knockdown reduced PGC-1α expression by ∼60% and increased pathological gene expression. To model exercise-induced physiological hypertrophy of cardiomyocytes, we stimulated the NRVMs with insulin-like growth factor 1 (IGF1). This is a stimulator of phosphoinositol-3-kinase/Akt-mediated growth pathways in cardiomyocytes that are observed with endurance training. It also reduces stress-related pathologic gene expression in cardiomyocytes^26,27^. IGF1 failed to reduce stress-related gene expression with PGC-1α knockdown (**Figure 2J**). Furthermore, intracellular ATP content increased with IGF1 and PGC-1α overexpression but the combination of PGC-1α silencing and IGF1 reduced ATP content by ∼80% relative to control siRNA treated cardiomyocytes (p<0.0001, **Figure 2K**) consistent with metabolic failure in the context of PGC-1α loss-of-function and exercise induced energy stress.

### Cardiomyocyte atrophy and increased SASP after exercise training in PGC-1α-deficiency

A striking feature of histological staining of the hearts of the KO_Ex mice was the relative loss of cardiomyocytes. PCM1 staining (nuclear marker of myocytes) showed a 25% reduction in PCM1+/DAPI+ cells suggesting a loss of cardiomyocytes (p<0.001, **Figure 3A**). Staining of a proliferation marker Ki67 also showed decreased Ki67+ cardiomyocytes in the KO_Ex hearts (67% decrease in KO_Ex vs. WT_Ex, p<0.0001, **Figure 3B**) with positive staining of noncardiomyocyte cells. KO_Ex hearts also demonstrated smaller cardiomyocytes by wheat-germ agglutinin (WGA) staining relative to WT_Ex counterparts (41% decrease, p<0.0001, **Figure 3C**). The findings of loss of cardiomyocytes as well as reduced cardiomyocyte proliferation and size suggested that PGC-1α was critical to protecting cardiomyocytes from atrophy in response to energetic stress related to exercise. While IGF1 promoted increased cardiomyocyte size as measured by sarcomeric α-actinin staining of treated NRVMs in vitro, it failed to increase cardiomyocyte size in NRVMs treated with PGC-1α siRNA relative to cells treated with PGC-1α siRNA alone (p<0.0001, **Figure S4A**).

**Figure 3:**
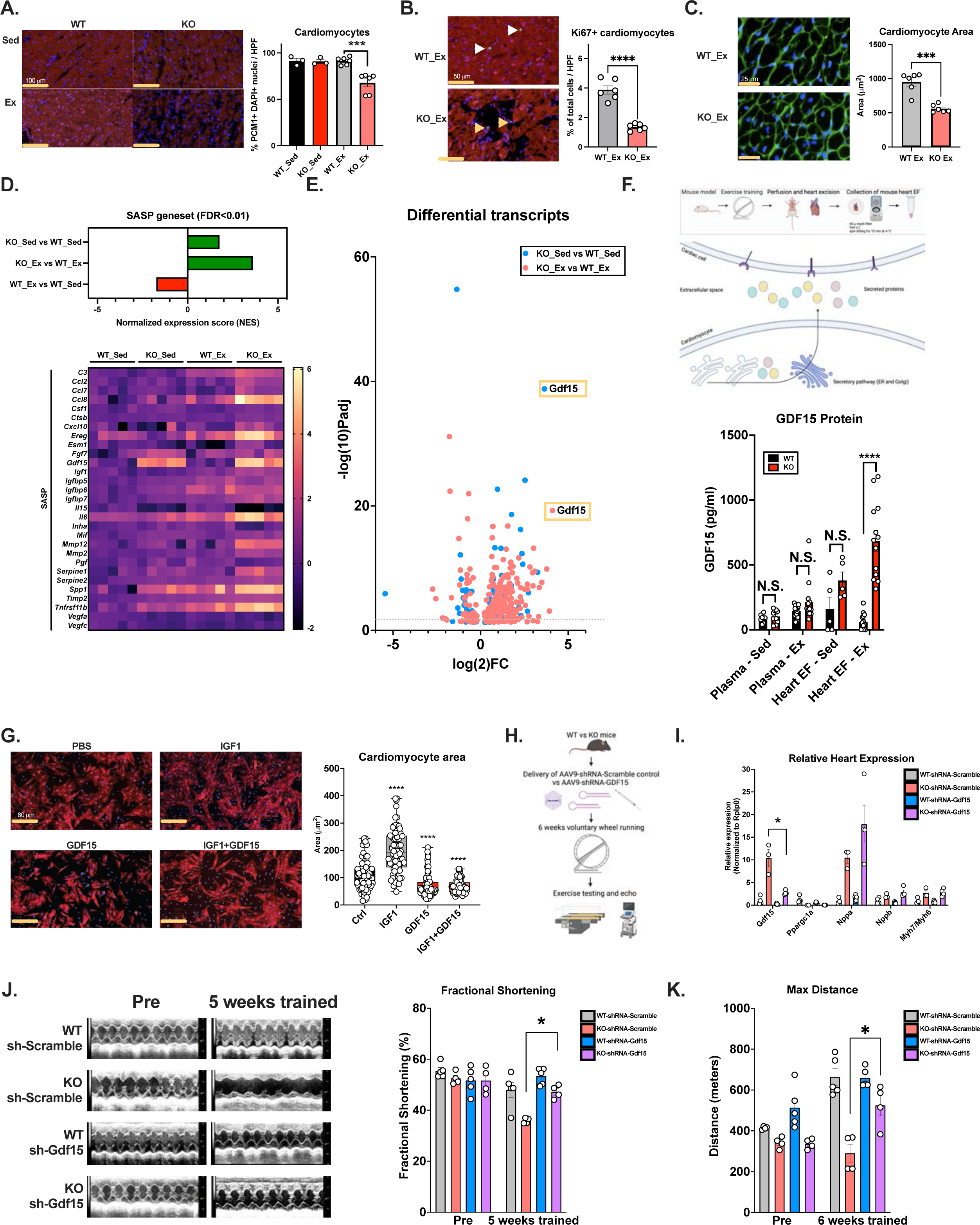
Cardiac atrophy, upregulated SASP and GDF15 in cardiomyocyte PGC-1α deficiency. **A**. Representative images of staining of frozen heart sections from indicated mice for PCM1 and DAPI (left) and quantification of PCM1+/DAPI+ staining per HPF (right). B. Representative images (left) and quantification (right) of Ki67 proliferation marker staining in WT_Ex and KO_Ex hearts. White arrowhead indicates Ki67+/PCM+/DAPI+ nuclei (cardiomyocytes). Yellow arrowhead indicates Ki67+/PCM-/DAPI+ nuclei (noncardiomyocytes). **C**. Wheat germ agglutinin (WGA) and DAPI staining of the hearts from B. **D**. (Top) Normalized enrichment scores for gene set enrichment for the SASP geneset (SenMayo) in indicated groups. For all comparisons FDR p-value < 0.05. (Bottom) Heat map of relative expression of SASP genes in RNA seq data expressed as log_2_(fold change) relative to the mean of the WT_Sed group. **E**. Volcano plot of differentially expressed protein-coding genes (FDR p-value < 0.05) in transcriptome from bulk heart RNA seq data for KO_Sed vs WT_Sed (blue) and KO_Ex vs WT_Ex (pink). **F**. (Top) Schematic diagram of extracellular space and approach to collection of heart extracellular fluid (EF). (Bottom left) Plasma GDF15 concentration from indicated groups of sedentary or exercised WT and KO mice measured by mouse GDF15 ELISA. (Bottom right) Heart EF GDF15 concentration using the same ELISA. **G**. (Left) Representative images for sarcomeric α-actinin staining of cardiomyocyte size in NRVMs treated for 48 hrs in serum-free media with PBS, IGF1 (1 ng/ml), GDF15 (500 ng/ml), or both IGF1 and GDF15. Cells were counter-stained with DAPI. (Right) Quantification of relative surface area of NRVMs in each group. **H**. Experimental design of AAV9-shRNA Scramble vs AAV9-shRNA Gdf15 expression and exercise tolerance experiment in WT and KO mice. **I**. Relative heart expression of indicated genes in mice from experiment in H. **J**. (Left) M-mode murine echocardiographic images of mice from G at rest prior to and at 5 weeks of exercise training. (Right) Quantification of FS from echo images in I. **K**. Max running distance in treadmill exercise test from mice in G prior to and at 5 weeks of exercise training. Data is expressed as mean +/- S.E.M. For A-C, H, J, and K, *P<0.05, ***P<0.001, **** P<0.0001, student’s unpaired T-test for the indicated groups. For F, ****P<0.0001, one-way ANOVA with Holm-Šídák’s multiple comparisons test to the Ctrl group.

To identify the mechanisms of atrophy in the KO_Ex hearts, we further studied their transcriptome. GSEA analysis of cell type signatures in sedentary and exercised KO vs WT mice also revealed that hearts from the KO mice demonstrated increased expression of genes related to aged cardiac cells (**Figure S4B**). Consistent with this, when we annotated all differentially expressed genes in the bulk sequenced transcriptomes and filtered by protein-coding genes encoding secreted proteins, there was an upregulation of genes constituting a program of negative regulation of tissue or cell growth (**Figure S4C**). Additionally, KO hearts demonstrated enriched expression of genes related to DNA damage-induced senescence, a finding that was weakly present even before exercise training (**Figure S4D**).

We found the KO hearts also enriched for the expression of senescence associated secretory phenotype genes^28^ (SASP; **Figure 3D**), a set of genes encoding secreted proteins that negatively regulate cell growth through paracrine and endocrine means in response to aging and stress^29–32^. Notably, KO_Sed mice already exhibited an upregulation of the SASP program (**Figure 3D**). We investigated individual genes that were upregulated in KO_Sed and KO_Ex hearts relative to their WT counterparts that comprised the SASP geneset (**Figure 3D**) and cell growth-related gene sets (**Figure S4C**). Among many known genes, one that was markedly upregulated (∼13-fold) in the sedentary KO mice was growth differentiation factor 15 (GDF15; **Figure 3E**). GDF15 was indeed the most enriched gene in the entire differential transcriptomes of the KO_Sed vs WT_Sed mice and the KO_Ex vs WT_Ex mice. *Gdf15* expression increased with PGC-1α siRNA treatment in NRVMs (**Figure S4E**). Conversely, muscle PGC-1α-transgenic mice overexpressing PGC-1α ∼8-10 fold in skeletal muscle and ∼3-4 fold in heart demonstrated a ∼50% downregulation of heart Gdf15 gene expression (**Figure S4F**).

### Heart-derived GDF15 limits cardiac function in response to exercise training in cardiomyocyte PGC-1α deficiency

Circulating GDF15 promotes energy expenditure in response to stress in several metabolic tissues through a central axis involving its receptor in the area postrema GFRAL^33,34^. To determine whether increased heart GDF15 expression in our model affected circulating levels, we measured plasma GDF15 using an ELISA. We found no difference in plasma GDF15 levels in sedentary, acutely-exercised or exercise trained WT vs. KO mice (**Figure 3F**). We next adopted a method previously developed to isolate muscle and fat extracellular fluid (EF) enriched for secreted myokines to the hearts of our mice^35^ (**Figure 3F**). We observed a ∼4.8-fold increase in GDF15 in heart EF of KO_Ex mice (P<0.0001, **Figure 3F**), mirroring the increase in Gdf15 gene expression. This suggests an elevation of locally secreted but not systemic GDF15 from the heart with PGC-1α cardiomyocyte deficiency.

Aside from its systemic role, heart-derived GDF15 limits pathological cardiac hypertrophy in response to pressure overload^36^. We found that exogenous GDF15 prevented physiological cardiomyocyte hypertrophy in response to IGF1 in NRVMs in vitro (**Figure 3G**). We thus hypothesized that early GDF15 upregulation and local secretion limited cardiac adaptation to exercise training in KO mice and that silencing cardiomyocyte *Gdf15* may prevent exercise-induced cardiac dysfunction. To test this, we developed a tool to silence cardiomyocyte Gdf15 expression using an adeno-associated virus serotype 9 (AAV9) vector expressing a short hairpin RNA (shRNA) against Gdf15 to administer to mice in vivo via tail-vein injection. We expressed this AAV vector or one expressing a control scramble shRNA sequence in 8-week old WT or KO mice for one week and then subjected them to endurance training for 6 weeks. We measured contractile function and exercise capacity before and after exercise training (**Figure 3H**). AAV9-Gdf15 shRNA reduced *Gdf15* expression in KO mouse hearts by ∼78% relative to KOs treated with the scramble shRNA by the end of exercise training (**Figure 3I**). KO mice expressing scramble shRNA demonstrated a 33% reduction in FS consistent with training induced cardiomyopathy (**Figure 3J**). Importantly, AAV-shGdf15 treated KOs retained their FS after training (51% pre-training vs 47% after 5 weeks, p=0.18; **Figure 3J**). They also demonstrated preserved exercise tolerance measured as max distance run during a treadmill tolerance test (66% improvement, p<0.05, **Figure 3K**). These data suggest that heart-derived GDF15 is an important and targetable contributor to exercise-induced contractile dysfunction and limited exercise capacity in the absence of cardiomyocyte PGC-1α function.

### PPARGC1A and GDF15 in cardiomyopathies

The exercise-induced cardiomyocyte atrophy and heart failure in the absence of PGC-1α suggested that improvement of myocardial mitochondrial function with exercise is necessary to prevent a maladaptive response to energetic stress. We hypothesized that the relationship of PGC-1α to the negative regulation of pathological protein secretion expression in other models may exist since downregulation of PGC-1α has been seen in murine models of heart failure. We performed comparative transcriptomics of bulk heart RNA seq from our model of exercise-induced heart failure with PGC-1α deficiency to that of PGC-1α-deficient pregnant female murine hearts exhibiting peripartum cardiomyopathy and of WT mice after transverse aortic constriction (**Figure S5**). We specifically annotated protein-coding transcripts for secreted protein gene expression and found that among upregulated secreted protein transcripts, the 124 genes overlapping across all models encoded programs related to fibrosis and extracellular matrix deposition (**Figure S5B**). Interestingly, 65 genes upregulated in the two PGC-1α-deficient heart failure mouse models (exercise and peripartum cardiomyopathy^16^) were enriched in programs related to immune cell chemotaxis (**Figure S5B**). Several components of the SASP were common to all 3 heart failure models, including GDF15 which was among the most highly upregulated across the 3 models (**Figure S5C**).

We next sought to extend this observation to human heart failure. In humans, germline deleterious variation in the PPARGC1A gene is poorly tolerated such that predicted loss-of-function (pLOF) variants are subjected to negative selection (gnomAD v4.0.0 server). We thus explored the relationship of cell-type resolved PPARGC1A expression in the hearts of humans with cardiomyopathies to investigate the relationship of PGC-1α function during myocardial energy stress to the observations from our murine model. Single nucleus RNA sequencing of 16 nonfailing, 11 dilated cardiomyopathy (DCM), and 15 hypertrophic cardiomyopathy (HCM) hearts previously identified almost 600,000 nuclei^37^. Using this study, we found that total PPARGC1A in the heart was predominantly expressed in the 3 different cardiomyocyte populations (**Figure 4A**). PPARGC1A cardiomyocyte expression was >28-fold higher than the next most expressing cell type which was vascular smooth muscle (VSMC). Comparing nonfailing (NF), DCM and HCM, PPARGC1A expression in cardiomyocytes was significantly reduced in both disease states, with ∼32% reduction in HCM and ∼38% reduction in DCM (p<0.0001 for both compared to NF, Wilcoxon rank-sum test; **Figure 4B**). We correlated total cardiomyocyte PPARGC1A expression with LV mass measures, left ventricular ejection fraction (LVEF), cardiomyocyte and macrophage frequency (**Figure 4C**). We found that cardiomyocyte PPARGC1A expression was correlated inversely with LV mass (p<0.05) and positively with relative cardiomyocyte proportion in the heart (p<0.05). A nonsignificant negative correlation was seen with macrophage proportion. Among cardiomyocytes expressing both genes, PPARGC1A and GDF15 were anti-correlated (p<0.05; **Figure 4C**). Consistent with this, meta-analysis of bulk-tissue RNA seq^38^ across tissues demonstrated a strong inverse correlation of PPARGC1A expression and GDF15 expression in the heart and muscle with weaker overall correlation across all tissues (**Figure S6**). Our data describes PGC-1α as an important restraint against energy stress-induced GDF15 expression and cardiomyocyte atrophy in the heart (**Figure 4D**).

**Figure 4:**
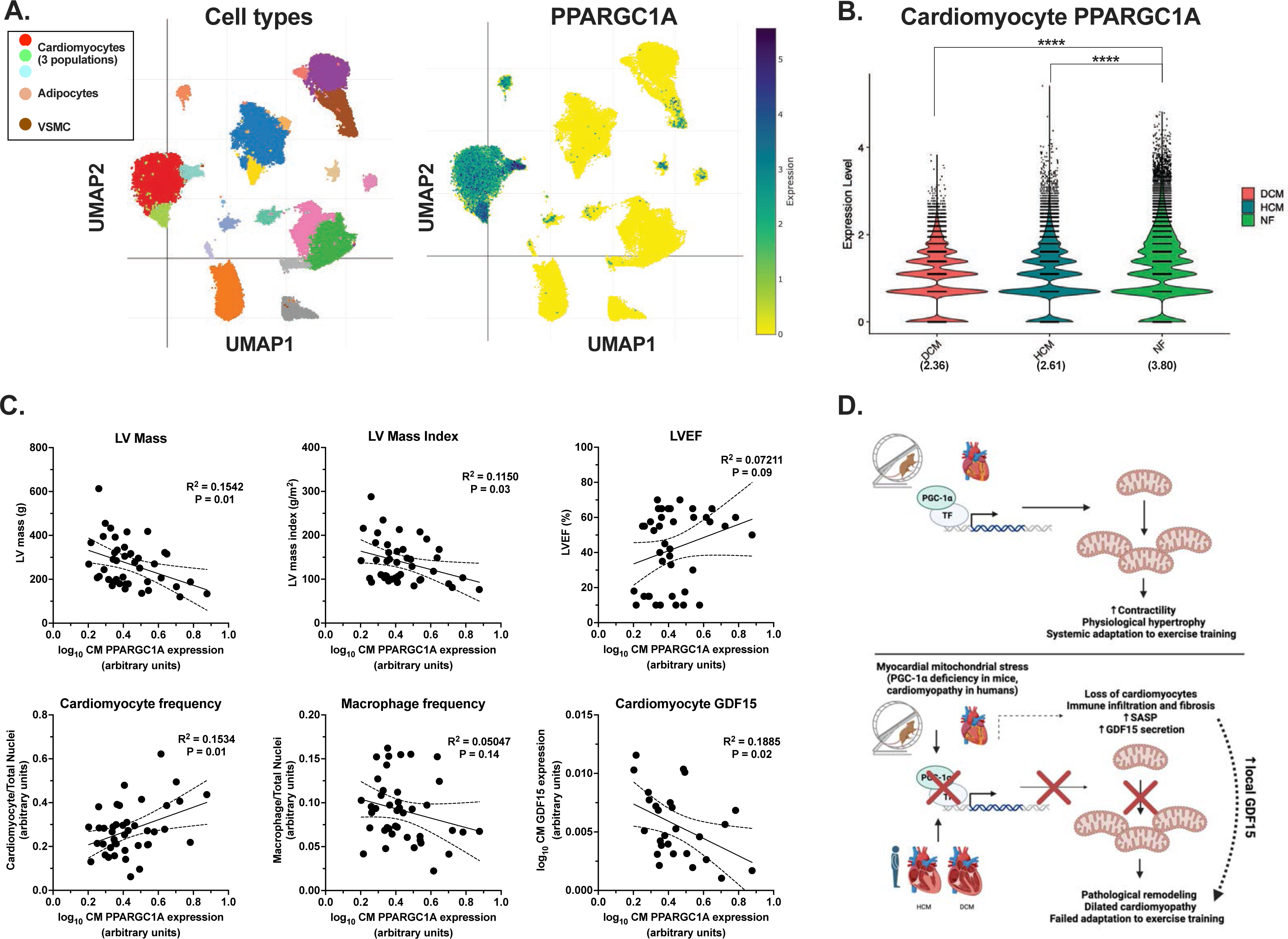
Cell-type resolved PPARGC1A expression in human cardiomyopathies. **A.** (Left) t-SNE plot showing 592,689 single cells from hearts from nonfailing human hearts (N=16), hearts from individuals with HCM (N=15) and hearts from individuals with DCM (N=11). (Right). Relative expression heatmap of PPARGC1A on the t-SNE plot. **B**. Relative expression of PPARGC1A in cardiomyocytes from A in NF, DCM and HCM hearts. Mean normalized expression is written below each group. **C**. Correlation between log(10) normalized PPARGC1A expression in cardiomyocytes (CM), to indicated measures with linear regression coefficient R2 and p-value indicated. **D**. Summary of physiological consequences of cardiomyocyte PGC-1α deficiency or relative reduction in the context of exercise and myocardial mitochondrial stress. For B, data is expressed as mean +/- S.E.M. ****P<0.0001, Wilcoxon Rank-sum test comparison to NF group.

## Discussion

Metabolic adaptation is a central component of tissue remodeling that confers many of the benefits of exercise^11^. PGC-1α is a single molecular entity that coordinates many of the adaptive programs in response to exercise in muscle. Here it does so through increasing mitochondrial numbers and function, inducing expression of genes related to fiber type switching and contractility, augmenting muscle innervation, myokine secretion and many other mechanisms^20^. PGC-1α increases with endurance training in the heart but the necessity of cardiomyocyte PGC-1α to exercise training is unknown. Moreover, the role of PGC-1α to established tenets of the endurance response from muscle (mitochondrial biogenesis, physiological hypertrophy, myokine secretion) in the heart has been unclear. Our study sought to address these and ascribe functions of PGC-1α to endurance training responses in the heart through studying genetic loss-of-function.

Our work includes several key new findings. First we show that cardiomyocyte PGC-1α is in fact required in mice for the cardiac adaptation to endurance exercise training. This is an important distinction from skeletal muscle, where some adaptation to exercise training occurs despite PGC-1α deficiency through increased mitochondrial function^13^. Unexpectedly we found that not only was cardiomyocyte PGC-1α required for a beneficial response to exercise, but its absence conferred the rapid onset of impaired resting contractility, immune fibrotic heart failure, and early systemic wasting (muscle and fat mass) after training. This establishes a distinct model of physiological stimulus-induced cardiomyopathy in the absence of heart PGC-1α separate from the placental-secreted anti-angiogenic vs pro-angiogenic factors that promote peripartum cardiomyopathy^16^. Here we show in male mice that an entirely separate mechanism of energetic stress via voluntary low-load exercise can promote similar dysfunction and upregulation of cardiac aging-related gene expression.

PGC-1α deficiency in the sedentary state itself promotes the SASP and an aging myocardial gene expression signature, suggesting that additional repeated energetic stresses (multiple pregnancies, endurance training) may provide a ‘second hit’ that confers cardiomyocyte atrophy in the absence of being able to enhance oxidative metabolism. Interestingly, we observe decreased cardiomyocyte proliferation markers and reduced cell size despite the known role of exercise in promoting these very features in the mouse heart and a marked transcriptional upregulation of the cell cycle. This suggests that states of increased cardiomyocyte senescence, such as aging, anthracycline chemotherapy treatment, and metabolic heart failure phenotypes such as diabetic and heart failure with preserved ejection fraction (HFpEF) may act through relative PGC-1α deficiency and that further energetic stress may be maladaptive.

Another key finding here is our evidence of the reciprocal relationship of PGC-1α to GDF15 expression in the heart. We found this initially in our animal model and show a directional relationship in mice, cardiomyocytes, and human transcriptomes. GDF15 has been an important developing therapeutic target for chemotherapy induced nausea and cancer cachexia particularly since the discovery of its hypothalamic receptor GFRAL^33,34,39^. It has also been shown that GDF15 expression increases in the heart as an anti-hypertrophic response during ischemic or pressure-overload related stress. Recently heart GDF15 was found to be maladaptively upregulated during doxorubicin treatment when combined with alternate-day fasting, a form of nutrient stress^40^. This work suggested that exogenous GDF15 was sufficient to confer cardiomyocyte atrophy in that model. Critically, we demonstrate by isolating the heart EF for comparison of GDF15 protein levels to plasma that heart-derived GDF15 causes a more profound elevation in local rather than systemic circulating GDF15. This is important in considering tissue-specific effects of modulating this circulating protein in the context of heart failure. We find that forced GDF15 reduction in the heart may be anti-atrophic and promotes exercise tolerance during energy stress. Our AAV silencing experiment provides proof-of-principle for heart-specific inhibition of local GDF15 to promote exercise and contractile function in states of relative PGC-1α deficiency and in cardiac cachexia.

Our work has important limitations which provide areas for further study. We predominantly assessed male mice here. A prior study of the cardiomyocyte PGC-1α-deficient mouse suggested an age-associated acceleration of excitation-contraction uncoupling in female mice as a cause of cardiac dysfunction in the sedentary state^41,42^. Our transcriptomic studies utilized bulk tissue sequencing to allow for harvesting prior to additional energetic stress that may be imparted by cardiomyocyte and immunocyte isolations. Future studies will compare transcriptomes using scRNA seq from these and related models to identify cell types of origin of the drivers of metabolic and immune-fibrotic dysfunction in addition to the SASP.

## Acknowledgements and Funding

This work was supported by National Institute of Health (NIH) grants R01 DK119117 to B.M.S., NIH grants R01 AG061034 and R35 HL155318 and American Heart Association (AHA) grant 23MERIT1038415 to A.R., a John S. LaDue Memorial Fellowship from Harvard Medical School to S.A.K., AHA grant 20CDA35310184 and NIH grant R21AG077040 to H.L., NIH grant K08HL140200 to J.R., a fellowship from Hope Funds for Cancer Research (HFCR-20-03-01-02) to H.-G.S., a Deutsche Forschungsgemeinschaft (DFG, German Research Foundation) grant (Projektnummer 461079553) to M.J.M., a NIH grant 1T32GM145407-01 to T.V., Damon Runyon Cancer Research Foundation Fellowship (DRG 120-17) and NIH grant K99 DK125722 to P.A.D., NIH grants HL163172 and K08 HL145019 to A.A., NIH grants K76AG064328 and R01HL170058 and the Yeatts Fund for Innovative Research to J.D.R.. The authors acknowledge Yoshiko Iwamoto (Center for Systems Biology, Massachusetts General Hospital) for help with sample staining, Jollanda Lako, Anuradha Kohli, and Jen Danielson for administrative support. The authors thank all members of the Spiegelman, Rosenzweig, Puigserver, and Chouchani labs for helpful discussions.

## Author Contributions

S.A.K.: Conceptualization, investigation, formal analysis, writing, funding. H.L.: Conceptualization, investigation. T.V.: Investigation, formal analysis. J.R.: Conceptualization. L.G.: Formal analysis. C.C.: Investigation. M.J.M.: Conceptualization, investigation. N.E.H.: Conceptualization. A.V.-C.: Investigation. A.L.S.: Investigation. J.L.: Investigation. C.C.: Investigation. H-G. S.: Conceptualization. K.A.B.: Investigation. A.K.: Investigation, formal analysis. R.F.: Investigation. D.B.: Investigation. P.T.E.: Resources. A.A.: Conceptualization, supervision. P.A.D.: Conceptualization, resources. P.P.: Conceptualization. J.D.R.: Conceptualization, supervision, resources. B.M.S.: Supervision, conceptualization, writing, resources, funding. A.R.: Supervision, conceptualization, writing, resources, funding.

## Materials and Methods

### Ethical approval

All mice were maintained and studied using protocols in accordance with the NIH Guide for the Care and Use of Laboratory Animals and approved by MGH Animal Care and Use Committees (protocol number 2015N000029) or by the Institutional Animal Care and Use Committee (IACUC) of Beth Israel Deaconess Medical Center (protocol number 072-2020).

### Animal studies

Mice with floxed alleles of Ppargc1a flanking exons 3 and 4 (JAX #009666), and mice containing the α-MHC-Cre transgene (B6.FVB-Tg(Myh6-cre)2182Mds/J, JAX #011038) were bred to generate mice in the indicated experimental groups. Genotyping for the Ppargc1a floxed allele and α-MHC-Cre allele was performed with Transnetyx (Cordova, TN). Wild type 12-week old male mice for the exercise timecourse experiment were obtained from The Jackson Laboratory (C57BL/6J, #000664). Hemizygous transgenic MCK-PGC-1α mice^14^ were bred in our animal facility on a C57BL/6J background. Wild-type littermates served as controls. Mice were fed a rodent chow diet with 12 hour light and dark cycles. Animal maintenance, exercise testing, endurance training and echocardiography were performed at the Massachusetts General Hospital Cardiovascular Research Center for Figure 1, and at the Beth Israel Deaconess Medical Center - Center for Life Sciences Small Animal Facility for Figure 3G-K.

Cardiac murine echocardiography was performed on unanesthetized mice. For experiments corresponding to Figure 1, a Vivid E90 cardiac ultrasound system (GE Healthcare) using an L8-l8i-D transducer. For experiments corresponding to Figure 3G-K, a Vevo 2100 microultrasound imaging system (VisualSonics, Toronto, Canada) was used. The heart was first visualized in long and short axis views followed by M-mode visualization of the short axis. Images were analyzed using EchoPACS software (Version 201, GE Healthcare). Parasternal short-axis M-mode images at the level of the papillary muscle were acquired at 10 mm depth to measure mid left ventricular dimensions at end-diastole (LV internal diameter at end diastole - LVIDd) and end-systole (LVESd). Interventricular septum (IVS) and LV posterior wall thickness (LVPW) dimensions were measured at end-diastole. Heart rate (HR) and fractional shortening (FS) was averaged from three consecutive beats. 3 measurements were obtained and averaged for each reported data point. Peak stress echo images were obtained immediately following acute exhaustive treadmill exercise protocol completion. Contractile reserve was measured as the difference between FS at peak stress and FS at rest.

For endurance exercise training, 12-week old male mice were individually housed in pre-autoclaved plexiglass cages containing a stainless steel running wheel (Mini-mitter, Starr Life Science, USA; diameter 11.4 cm) containing a tachometer. Mice were allowed to run voluntarily in continuity for the duration of training (6 weeks for acute exhaustive exercise testing and peak-stress and rest echocardiography, 8 weeks till euthanization and collection of tissues for molecular analyses). Mouse wheel running activity was recorded for at least 3 weeks of the running period (weeks 2-5 since initiation).

For acute treadmill exhaustive exercise testing, for 3 days prior to planned formal exhaustive protocol exercise testing, mice were acclimated to the treadmill. For experiments corresponding to Figure 1 at MGH an automated treadmill was used (Columbus Instruments). For experiments corresponding to Figure 3G-K at BIDMC (Columbus Instruments) a non-automated treadmill was used. For acclimatization, for 3 consecutive days mice were subjected to walking at a pace of 5 meters/min for 15 min (day 1), 5-10 meters/min for 15 min (day 2), and 5-30 meters/min for 15 min (day 3) with the treadmill incline set to 10°. For the acute exhaustive running protocol in experiments corresponding to Figure 1, a warmup period lasted for 5 min and then the treadmill was accelerated to require a mouse’s power output to increase by 3mWatt/min (from a starting power of 10mWatts) until it reached exhaustion; this typically corresponded to a treadmill acceleration of 1.5-2m/min^2. For experiments corresponding to Figure 3G-K, a warmup period of 10 min at 12 meters/min was followed by stepwise increase in speed by 2 meters/min every 5 minutes. Mice ran until exhaustion was reached which was determined to be that the mice could not keep pace with the treadmill for 3 seconds without falling back onto the resting pattern and that this behavior repeated 3 consecutive times. At that point the mouse was removed from the treadmill. For Figure 1, power and work were calculated based on the distance run, the angle of the treadmill, the weight of the mouse, and the velocity achieved.

For tissue analyses, mice were euthanized with 4% isoflurane inhalation followed by cardiac puncture and exsanguination. Tissues were snap frozen in liquid nitrogen. Apical sections of the heart (approximately 15-20 mg) were cut using a sterile razor using a glass cutting block and quickly frozen in OCT preservative (Sakura) in plastic preservative blocks using 2-methylbutane and dry ice.

For heart extracellular fluid (EF) collection, mice were administered continuous 4% isoflurane by inhalation, the thorax was opened, an incision was made in the right atrium, a 30 gauge needle was injected in the left ventricular apex and connected to a digital peristaltic pump (Reglo, Harvard Bioscience Inc.). 10 cc of cold PBS was infused at a rate of 10 cc/min into each heart, followed by rapid excision. Hearts were dried for 5 seconds on a Kimwipe (Millipore Sigma). Then following the protocol of Mittenbühler et al., the heart was placed in a 20 µm nylon mesh filter (Millipore Sigma), folded in half twice, and placed in a 2 ml Eppendorf tube and spun in a microcentrifuge for 10 minutes at 4 °C. The fluid spun through the folded mesh filter was defined as heart EF and frozen at -80 °C. For GDF15 ELISA, the R&D DuoSet ELISA for mouse GDF15 was used according to the manufacturer’s instructions. For plasma, 10 µl per sample was used. For heart EF, 5 µl per sample was used. Each sample was run in duplicate.

### Adeno-associated virus

Recombinant adeno-associated virus serotype 9 (AAV9) vectors were cloned and propagated by VectorBuilder (VectorBuilder Inc, Chicago, USA). Both AAV9-shRNA-Gdf15 and AAV9-shRNA-Scramble vector plasmids harbored shRNA expression driven by the U6 promoter and CMV promoter driven eGFP expression downstream of the shRNA insert. Vector plasmids utilized are the following: AAV9-shRNA-Gdf15 - VB900139-2079dda, AAV9-shRNA-Scramble - VB010000-0023jze. Recombinant AAV was produced in HEK293T cells. AAVs were administered at a dose of 5x10^11^ viral genomes (v.g.)/mouse by diluting in a final volume of 200 µl PBS via tail-vein injection using 1 ml 30 Ga insulin syringes (B.H. Supplies). Mice were maintained in the sedentary state for 1 week following AAV administration and then treadmill testing, echocardiography and exercise training was initiated.

### Quantitative real-time PCR for mRNA expression

Total RNA from each tissue or frozen cells was extracted using TRIzol (Invitrogen), purified with RNeasy Mini spin columns (Qiagen), and reverse transcribed using a HighCapacity cDNA Reverse Transcription kit (Applied Biosystems). The resulting cDNA was analyzed by RT-qPCR using SYBR green fluorescent dye 2x qPCR master mix (Promega) in a QuantStudio 6 Flex Real-Time PCR System (Applied Biosystems). The Rplp0 mRNA was used as a loading control, and fold change was calculated using the ΔΔCt method. Primer sequences were generated using the IDT PrimerQuest tool (Integrated DNA Technologies).

### RNA sequencing

RNA sequencing was performed by Novogene. Libraries were constructed from polyA-selected RNA using a NEBNext Ultra Directional RNA Library Prep Kit (New England Biolabs) and sequenced on Illumina HiSeq2500 instrument. The R package DESeq2 was used for differential gene expression analysis. Genes were considered differentially expressed if upregulated by log2FC>1 or downregulated by log2FC<−1 with an adjusted P-value <0.05 (using Benjamini-Hochberg correction). Gene Ontology (GO) overrepresentation analysis was performed using the database using the WebGestalt program^24^. For all differentially expressed genes, a metric was computed as the product of log(2)FC and −log10(p-value). Gene set enrichment analysis (GSEA^21,22^) using the ‘Classic’ mode was used to calculate enrichment scores and statistics using DESeq2 normalized expression levels after log(2)FC ranked expression by gene. Genes for which average DESeq2 expression values were 0-5 were excluded for GO and GSEA analysis pre-ranked gene lists. For GSEA analyses, Hallmark gene sets (MsigDB, MH) and cell type signature gene sets (MsigDB, M8) were used. For senescence geneset analysis, MsigDB GO genesets were queried. For the SASP gene set analysis, a consensus senescence associated secretory phenotype SASP gene set (SenMayo) was used.

### Immunoblotting

For immunoblotting of murine tissues, approximately 40-50 mg of each tissue was used. Tissues were prepared in ice-cold RIPA buffer (150 mM NaCl, 1% Nonidet P-40, 0.5% sodium deoxycholate, 0.1% SDS, 50 mM Tris pH 7.4) supplemented with Complete EDTA-free protease inhibitor [Roche]). Tissues were homogenized using a Polytron PT 10-35GT homogenizer (Kinematica). Protein concentration of the lysates was determined by bicinchoninic acid assay (Pierce), followed by denaturation in Laemmli buffer (50 mM Tris pH 6.8, 2% SDS, 10% glycerol, 100 mM DTT, and 0.05% bromophenol blue). Proteins were resolved by SDS-PAGE in 4-12% NuPAGE Bis-Tris gels (Invitrogen) and transferred to polyvinylidene difluoride membrane with 0.45 μm pore size (ImmobilonP). Membranes were blocked with Tris-buffered saline with 0.05% Tween-20 (TBST) containing 5% dried nonfat milk (Biorad). Primary antibodies were diluted in TBS-T containing 5% dry nonfat milk (Biorad). Membranes were incubated overnight at 4 °C with primary antibody. For secondary antibody incubation, anti-rabbit HRP (Promega) or anti-mouse HRP (Promega) was diluted in TBST containing 5% dried nonfat milk. Secondary antibodies were visualized using enhanced chemiluminescence western blotting substrates (Pierce), Immobilon Crescendo HRP substrate (Millipore). Primary antibodies in this study include the following: Col3a1 (Novus Biologicals, NB600-594), periostin (Abcam, ab92460), Vimentin (Cell Signaling Technology, 5741S), smooth muscle actin (SMA) (Thermo Fisher, NBP233006A) TGFβ1 (Cell Signaling Technology, 3711S), GAPDH (Cell Signaling Technology, 97166S). Secondary antibody used in this study was the following: Anti-Rabbit IgG (H+L), HRP Conjugate, (Promega, W4011).

### Neonatal rat ventricular myocyte isolation and experiments

Primary neonatal rat ventricular cardiomyocytes (NRVMs) were isolated as described previously^25,43^. Isolated NRVMs were purified by pre-plating and percoll gradient centrifugation. NRVMs were plated in 6-well plates precoated with gelatin (Sigma) at 0.8x10^5^ cells per well and cultured in NRVM medium (DMEM supplemented with 5% FBS and 10% horse serum) for 24 hours. Before treatment, NRVMs were synchronized and cultured in DMEM containing 0.2% FBS. Twenty-four hours after plating, cells were treated with adenovirus expressing PGC-1α as described previously^35^ (at multiplicity of infection of 100), or transfected with control siRNA (Thermo, 4390843) or sirRNA to rat Ppargc1a (s135986). Transfections were performed using RNAiMax transfection reagent (Thermo) according to the manufacturer’s protocol in Optimem medium containing 5% FBS with a final siRNA concentration of 25 nM. 18 hours later media containing siRNA or adenovirus was removed and replaced with NRVM medium. Then 6 hours later NRVMs were treated with PBS or 100ng/mL IGF1 (291-G1-01M; R&D Systems) to elicit physiological hypertrophy for 48 hours prior to cell harvesting for experiments. For GDF15 treatment of NRVMs, NRVMs were plated as above in 6-well plates coated with gelatin at a density of 600,000 cells/well in NRVM medium. 48 hours later, cells were incubated in serum free DMEM containing 1 mM sodium pyruvate (Gibco) containing PBS, IGF1 (1 ng/ml), recombinant human GDF15 (500 ng/ml; 957GD025CF), or both IGF1 and GDF15 in a total volume of 2 ml/well at the indicated concentrations for 48 hrs.

For quantitative RT-PCR, RNA from each well was harvested in 1 ml of TRIzol. ATP concentration was quantified using the ATP determination assay from Thermo (A22066). Cells were counted on a light microscope (at least 10 high power fields per well were counted) and used for normalization, and cells were lysed in 1x Cell Lysis Buffer (Cell Signaling Technology, 9803) supplemented with Complete EDTA-free protease inhibitor [Roche] before performing the ATP determination assay.

For measurement of NRVM cell size, cells were washed and fixed in 4% paraformaldyehyde for 15 min, then washed with PBS. Regions of interest for staining were outlined with ImmEdge pen and then blocked for 30 min at room temperature using 4% normal goat serum, 1% BSA, 0.2% triton-X 100 in PBS. After rinsing the plates, cells were stained with anti-sarcomeric alpha actinin antibody (Abcam, ab9465) 1:200 in antibody buffer (1% BSA, 0.1% triton-X 100 in PBS) for 2 hours a room temperature. Plates were washed and stained with goat-anti-mouse Alexa 594 antibody (Invitrogen A11032) 1:500 in antibody buffer for 2 hours at room temperature and then washed and stained with DAPI and then imaged with Leica DM500B Microscope. Cardiomyocyte cross-sectional area (∼100 cells per well) was measured from randomly selected sections per heart using ImageJ (NIH).

### Immunohistochemistry of mouse heart sections

Apical sections were stained with Wheat Germ Agglutinin (WGA), Alexa FluorTM 594 (W11262, Thermo Fisher Scientific) for cell size measurement. WGA stained slides were scanned by a digital slide scanner, NanoZoomer 2.0-RS (Hamamatsu, Japan). For cardiomyocyte analysis, immunofluorescent staining was performed. Anti-PCM1 antibody (HPA023374, SigmaAldrich, cardiomyocyte specific marker) was incubated at 4°C overnight and a biotinylated secondary antibody followed by streptavidin-DyLight 594 (BA1000 and SA-5594, Vector Laboratories) were used for cardiomyocyte identification. Nuclei were counterstained with DAPI (D21490, Thermo Fisher Scientific) and the slides were imaged on a Leica DM500B Microscope. Cardiomyocyte cross-sectional area (∼100 cells per heart) was measured from six randomly selected sections per heart using ImageJ from WGA stained cells (NIH). For cardiomyocyte proliferation marker quantification, anti-Ki67 antibody (clone: SolA15, 14-5698-82, Thermo Fisher Scientific) and Alexa Fluor 488 goat anti-rat IgG secondary antibody (A-11006, Thermo Fisher Scientific) were applied. For CD68 staining, BioLegend (valid) # 137001 ) sections were incubated with a rat anti-mouse CD68 antibody (BioLegend (valid) #137001) for 2 hr at room temperature. Alexa Fluor 568 goat anti-rat IgG antibody was used as a secondary antibody. For CD3 staining, anti-mouse CD3 primary antibody was used for incubation for 2 hrs at room temperature (BD Biosciences (valid) # 555273 ), followed by Alexa fluor 568 goat-anti-rat IgG secondary antibody. For Picrosirius Red Staining we followed the manufacturer’s instructions (Polysciences, #24901). Briefly, frozen OCT heart sections were stained, dehydrated, cleared, and mounted, followed by imaging using light microscopy. Quantification of positive staining per HPF was quantified in Image J (NIH).

### Single Nuclear RNA sequencing (snRNA-seq) analysis

Data for snRNA-seq was downloaded from The Broad Institute Single Cell Portal^37^. The three cardiomyocyte clusters were subset from the entire dataset, normalized using SCTransform and analyzed using a Seurat pipeline in R 4.2.1. Correlations of cardiomyocyte gene expression with immune cell proportions and clinic measures of heart function from Chaffin, et al, *Nature*, 2022^37^ were performed in Prism GraphPad 10.1.0.

### Bulk RNA sequencing immune deconvolution

FPKM counts from the bulk heart RNA sequencing previously described was loaded into R 4.2.1. The immunedeconv R package (https://doi.org/10.1093/bioinformatics/btz363) was then used to deconvolute the immune signatures in the RNA sequencing using mMCPCounter (https://doi.org/10.1186/s13073-020-00783-w) and DCQ (https://doi.org/10.1002/msb.134947) on the native mouse data. Relative proportions were graphed, and statistical tests were run in Prism GraphPad 10.1.0.

### Statistics, analysis and reproducibility

Replicate numbers are indicated in figure legends. Sample sizes were determined based on prior experiments using similar methods. Unless otherwise stated, data are presented as mean +/- standard error of the mean. Graphing and statistical analyses, including two-tailed Student’s t-test, one-way ANOVA, and Fisher’s LSD, were performed using GraphPad Prism 10 (GraphPad). Images in the figures were created using Biorender (Biorender.com).

---

**Figure S1:**
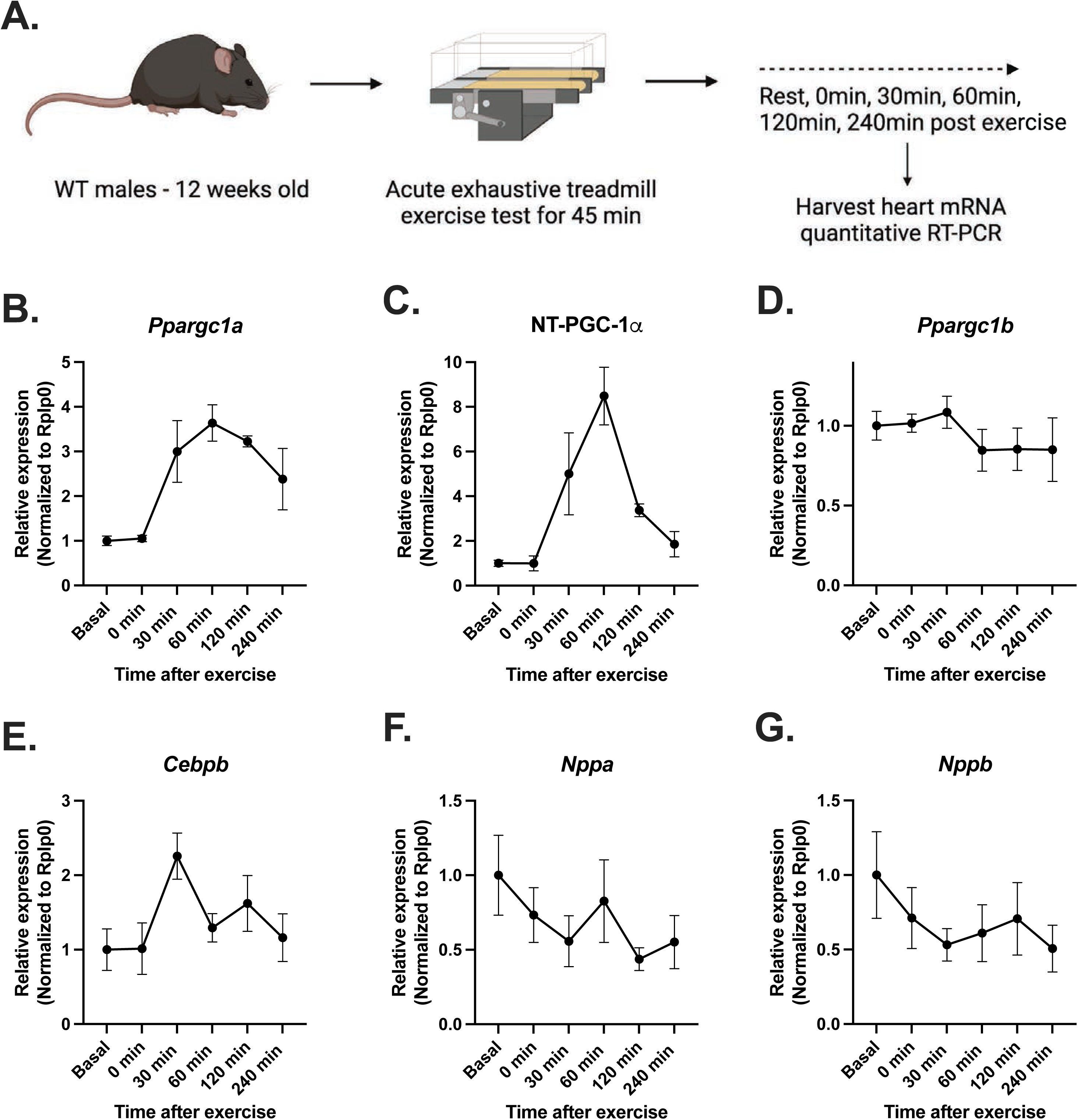
Acute treadmill exercise heart gene expression timecourse. **A.** Experimental design of timecourse. **B-G.** Relative mRNA levels of the indicated genes at the indicated timepoints following acute treadmill exercise for 45 min. N=6 mice per group. Data is presented as mean +/- S.E.M.

**Figure S2:**
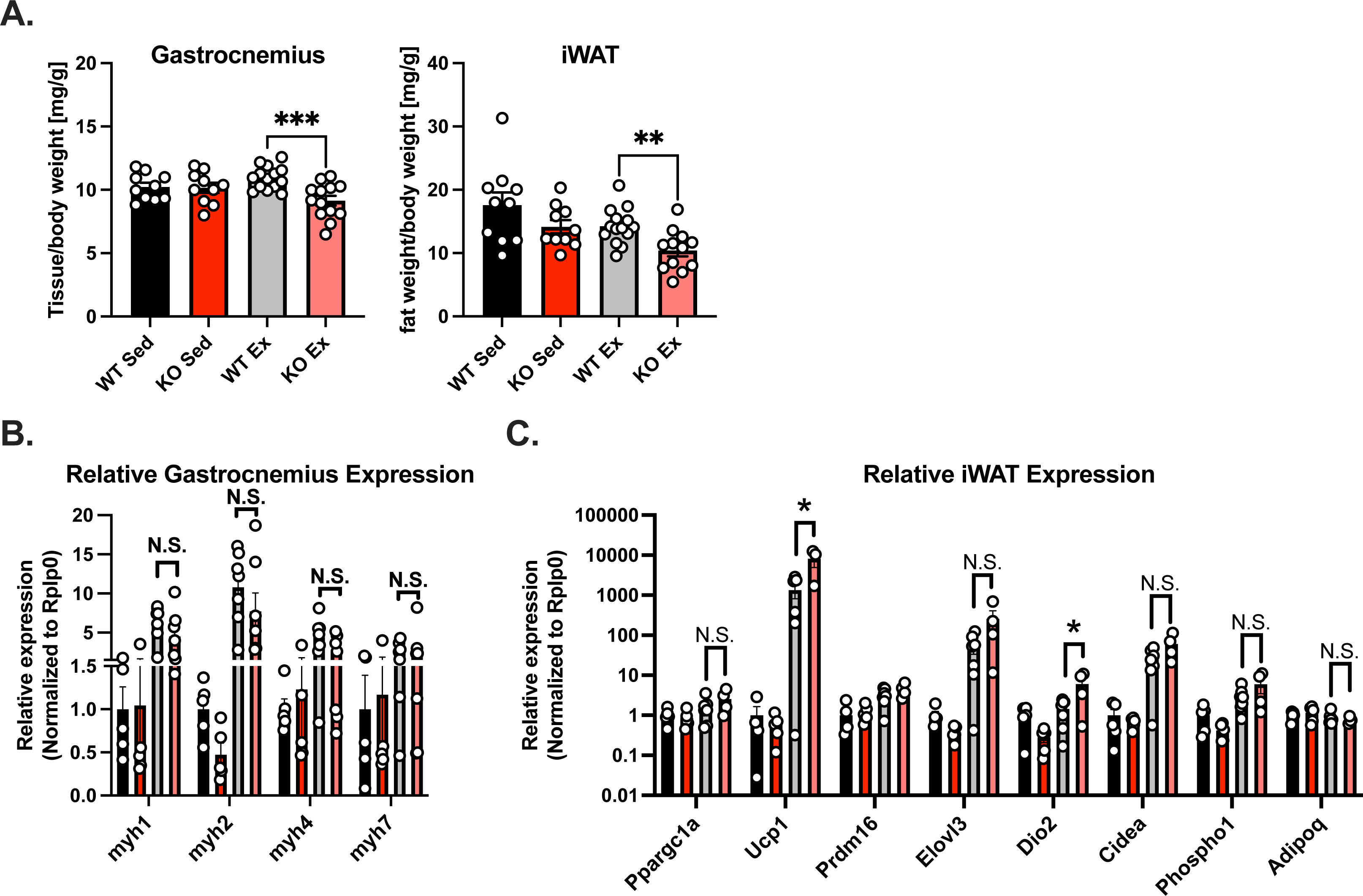
Fat and muscle mass and gene expression in inguinal white adipose tissue in sedentary vs exercise trained WT and KO mice. **A**. Relative inguinal white adipose tissue (iWAT) and gastrocnemius weights normalized to body weight in indicated mice. **B**. Relative expression of the indicated genes in iWAT normalized to that of Rplp0. **C**. Relative expression of indicated genes in gastrocnemius normalized to that of Rplp0. Data is expressed as mean +/- S.E.M.. *P<0.05, student’s unpaired T-test. Data is presented as mean +/- S.E.M.

**Figure S3:**
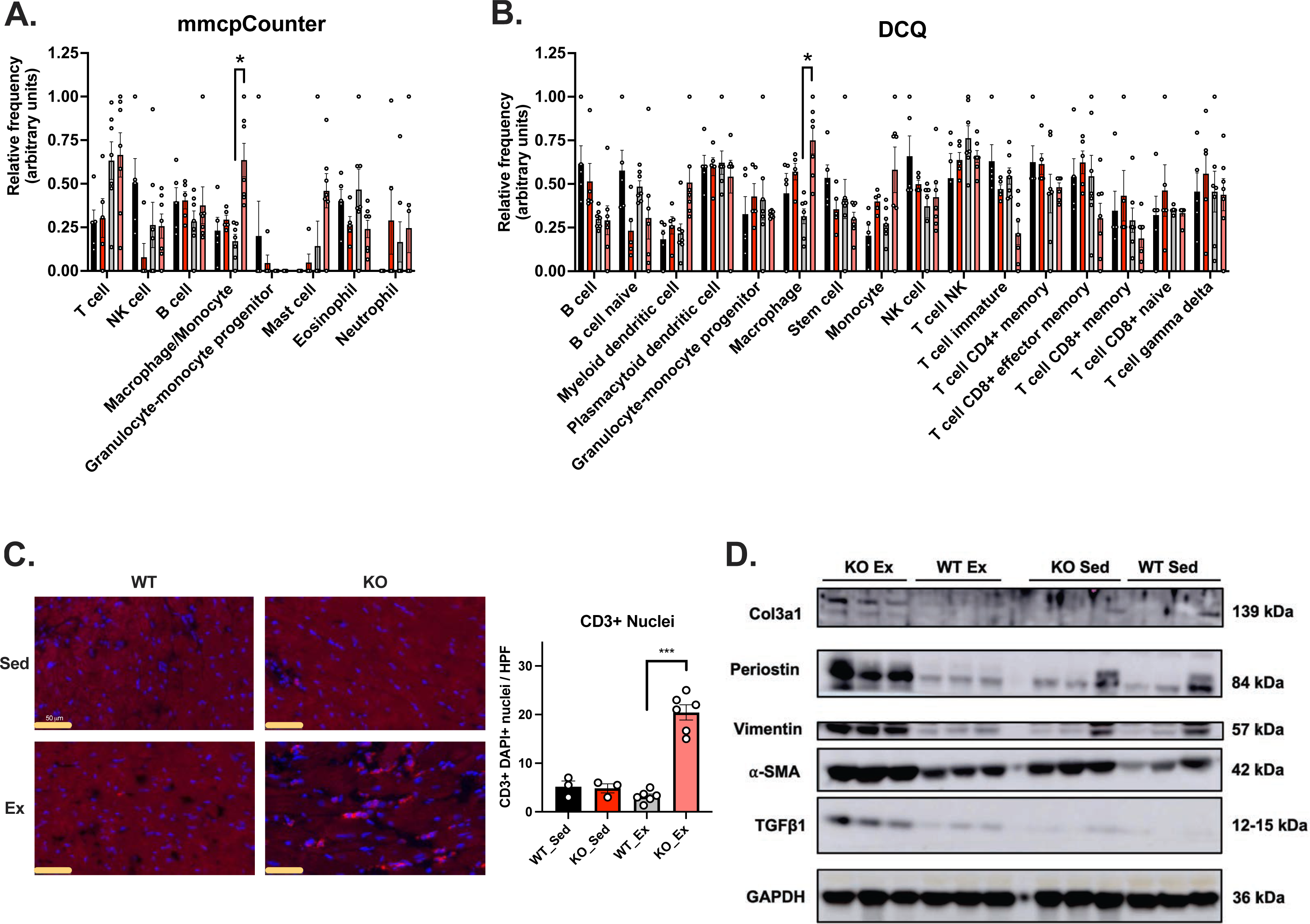
Relative immunocyte content in hearts of sedentary and exercise trained WT and KO mice. **A**. Immune cell relative frequency from deconvolution analysis of heart bulk RNA seq data corresponding to Figure 2 using the murine Microenvironment Cell Population counter (mMcp counter) tool. **B**. Immune cell deconvolution analysis of RNA seq data from A using the Digital Cell Quantification (DCQ) tool. **C**. Representative images of CD3 staining of frozen heart sections from mice in A (left) and quantification of CD3+/DAPI+ stains per HPF. **D**. Immunoblot for fibrosis markers from heart lysates of mice in A. N=3 per group. Data is expressed as mean +/- S.E.M.. *P<0.05, **P<0.01, ***P<0.001, **** P<0.0001, student’s unpaired T-test. Data is presented as mean +/- S.E.M.

**Figure S4:**
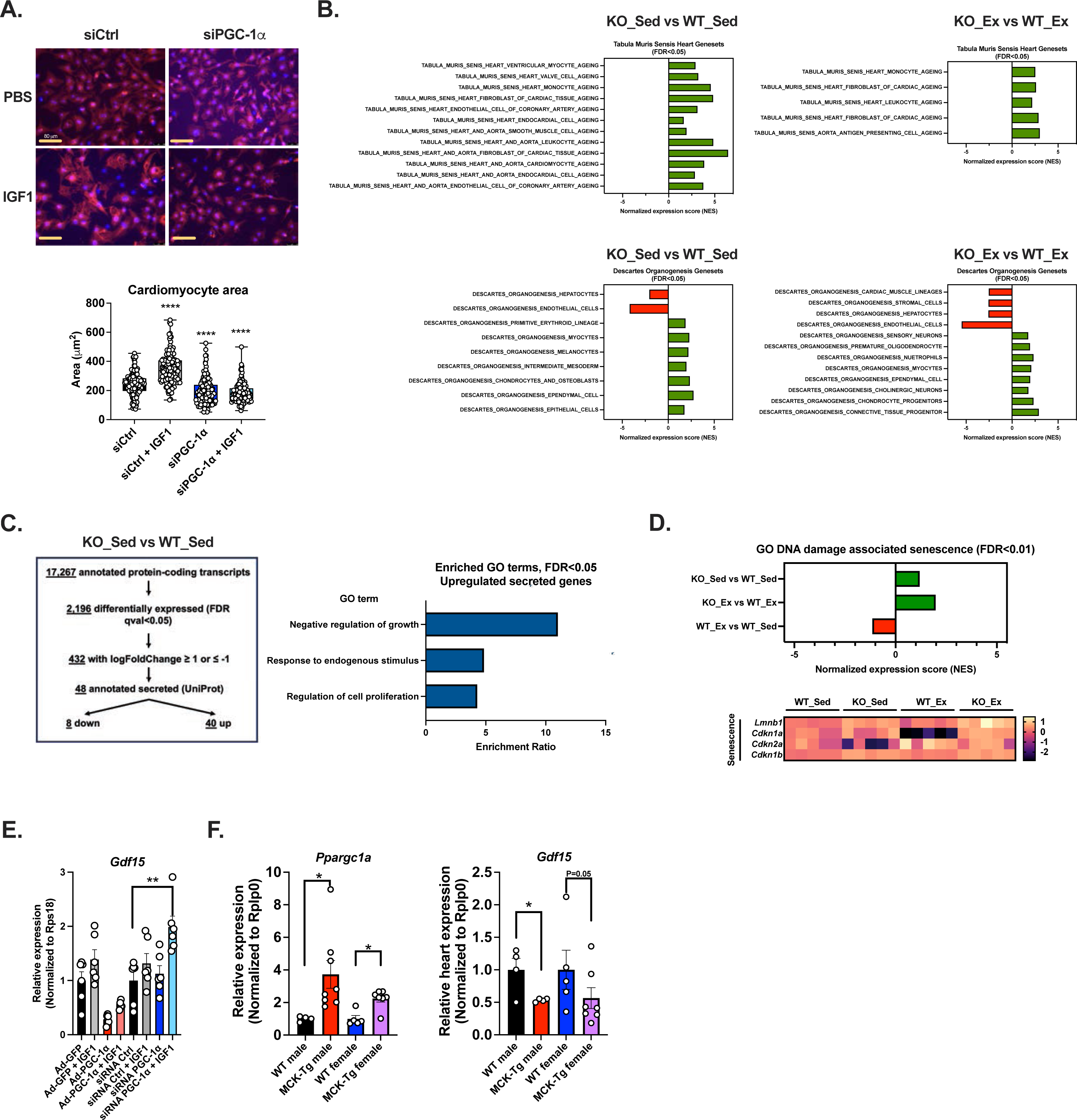
PGC-1α gain- or loss-of-function, cardiomyocyte size, and GDF15 in NRVMs and mice. **A**. (Top) Representative images from sarcomeric α-actinin staining of NRVMs after adenoviral PGC-1α overexpression or siRNA PGC-1α knockdown in the context of IGF1 stimulation of physiological hypertrophy as described in Figure 2J-K. Cells were costained with DAPI to identify nuclei. (Bottom) quantification of relative cell size from staining. **B**. Geneset enrichment scores for indicated group comparisons for Tabula Muris Sensis aging cell type gene signatures (top) and Descartes Organogenesis cell type gene signatures (bottom), both sets obtained from MsigDB (gsea-msigdb.org/gsea/msigdb/mouse/genesets.jsp) **C.** (Left) Approach to prioritizing genes encoding secreted proteins in bulk RNA seq analysis from mouse model in Figure 2. (Right) Enrichment scores for significantly enriched gene ontologies (FDR p-value < 0.05) for 40 genes corresponding to upregulated secreted proteins from prioritization in B. **D**. (Top) Geneset enrichment scores for MsigDB GO DNA damage associated senescence geneset. (Bottom) Heatmap plotting log(2)FC relative to the WT_Sed group for relative gene expression for indicated senescence gene markers. Relative expression of indicated genes in NRVMs according to experimental design in A. **E**. Relative heart expression of indicated genes in male and female muscle transgenic mice overexpressing PGC-1α. Data is expressed as mean +/- S.E.M.. *P<0.05, **P<0.01, ***P<0.001, **** P<0.0001, student’s unpaired T-test. Data is presented as mean +/- S.E.M.

**Figure S5:**
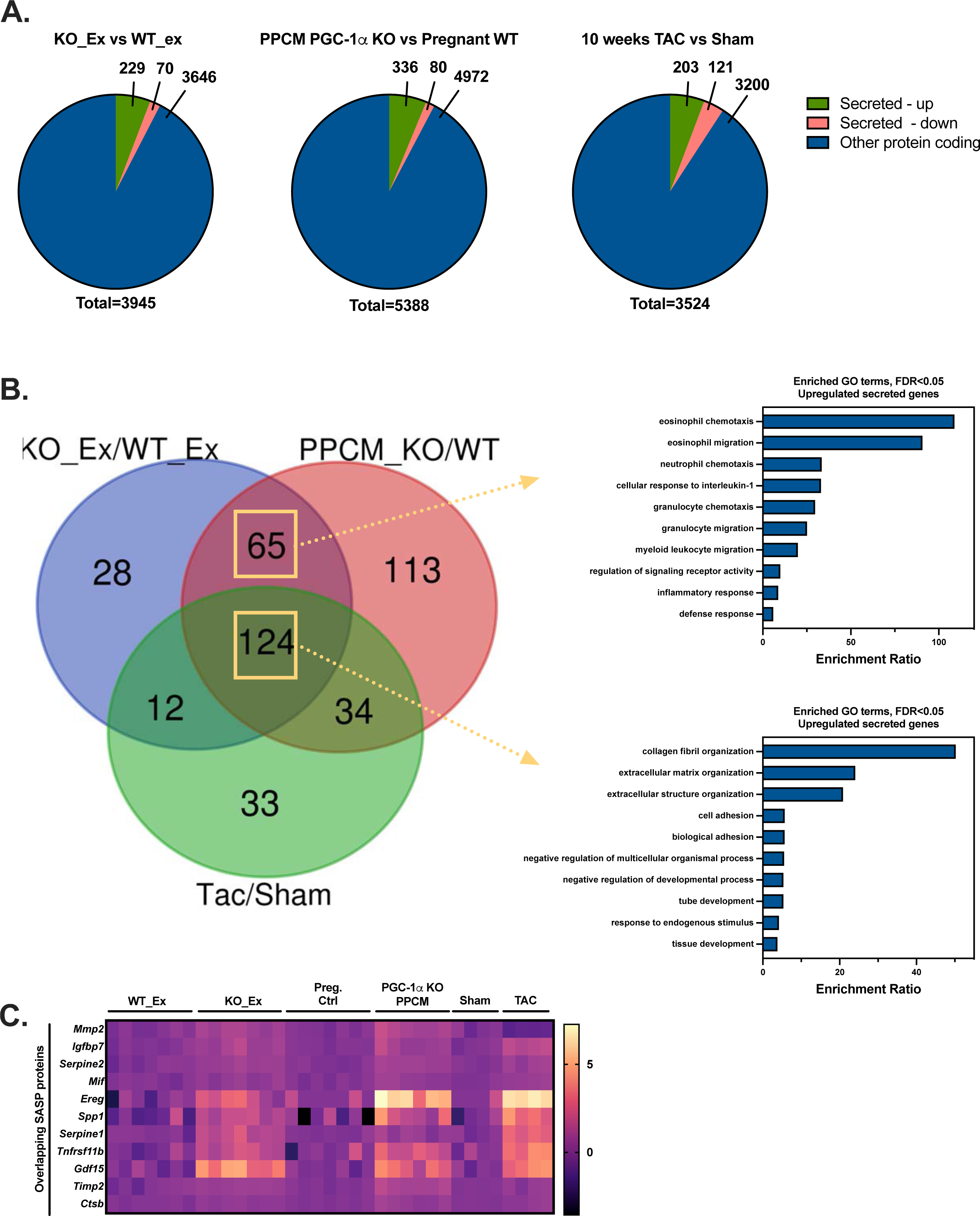
Secretory protein expression phenotype across murine heart failure models. **A.** Pie charts showing fraction of upregulated secreted protein, downregulated secreted protein and remaining protein coding transcripts from differential transcriptome gene expression analysis of bulk heart RNA seq from KO_Ex vs WT_Ex mice (N=7/group), PGC-1α WT pregnant female (N=7) vs PGC-1α cardiomyocyte KO female mice after 2 pregnancies (N=6), and WT mice that underwent sham (N=4) vs transverse aortic constriction surgeries (N=4). Cutoff for transcripts included for pairwise comparisons was FDR<0.05. Secreted protein transcripts were annotated using the UniProt database. **B**. (Left) Venn diagram of overlapping and distinct differential transcriptomes that were upregulated in the pairwise comparisons from A across the 3 comparisons. (Right) Arrows point to bar charts of enrichment scores for top 10 overrepresented GO terms for the gene sets indicated by the boxes from the Venn diagram. **C**. Heatmap plotting log(2)FC relative to the control groups for each of the 3 studies in A for the normalized expression of the genes corresponding to the SASP geneset.

**Figure S6:**
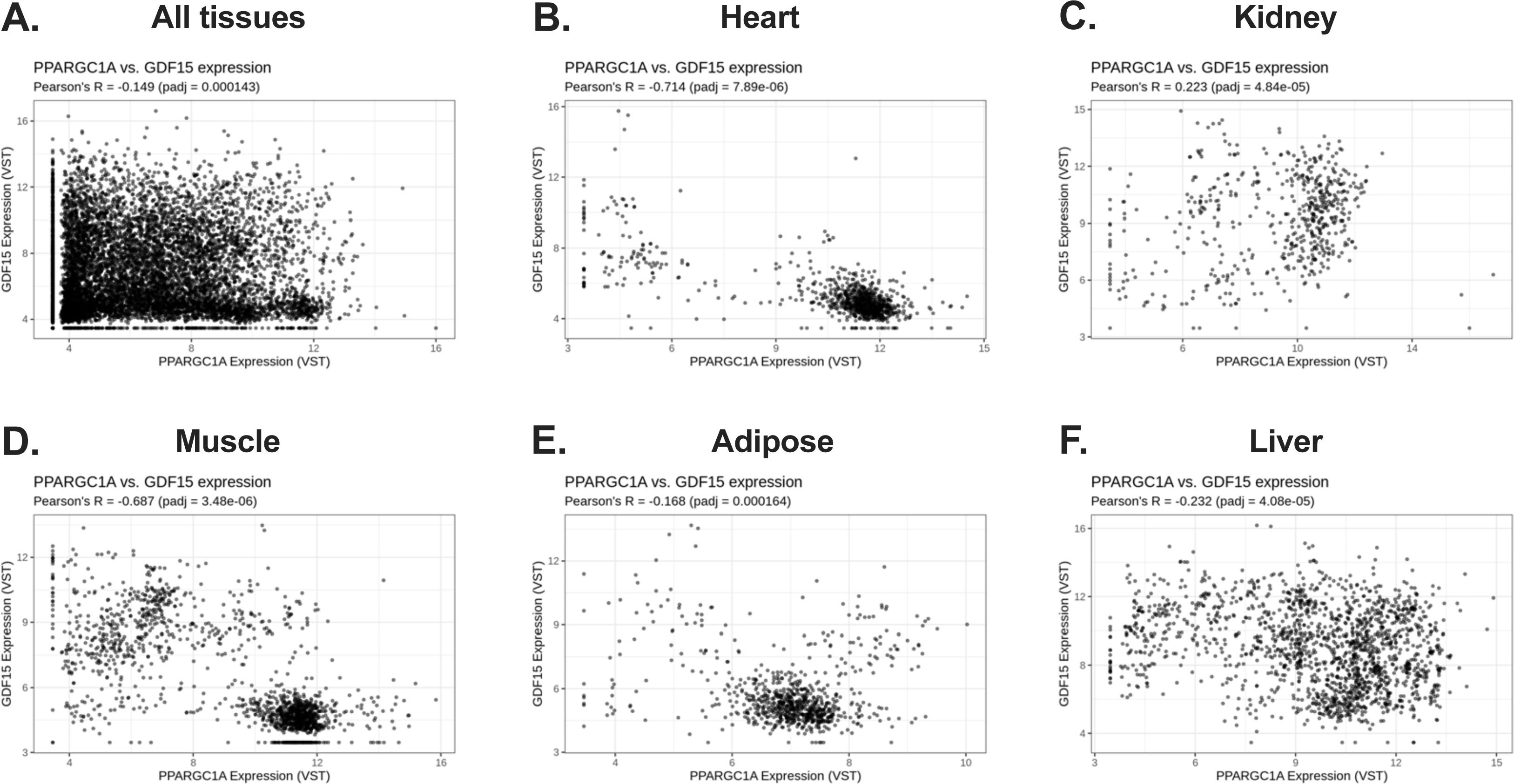
PPARGC1A correlations with GDF15 in human tissues: **A-F**. Correlation of PPARGC1A expression with GDF15 expression in indicated tissues from deposited bulk RNA seq data (NCBI GEO) using the ARCHS^4^ database. Correlation coefficient from linear regression and p-value are shown.

